# A mean-field theory for predicting single polymer collapse induced by neutral crowders

**DOI:** 10.1101/2023.07.12.548683

**Authors:** Quentin Chaboche, Gerardo Campos-Villalobos, Giuliana Giunta, Marjolein Dijkstra, Marco Cosentino-Lagomarsino, Vittore F. Scolari

## Abstract

Macromolecular crowding can induce the collapse of a single long polymer into a globular form due to depletion forces of entropic nature. This phenomenon has been shown to play a significant role in compacting the genome within the bacterium *Escherichia coli* into a well-defined region of the cell known as the nucleoid. Motivated by the biological significance of this process, numerous theoretical and computational studies have searched for the primary determinants of the behavior of polymer-crowder phases. However, our understanding of this process remains incomplete and there is debate on a quantitatively unified description. In particular, different simulation studies with explicit crowders have proposed different order parameters as potential predictors for the collapse transition. In this work, we present a comprehensive analysis of published simulation data obtained from different sources. Based on the common behavior we find in this data, we develop a unified phenomenological model that we show to be predictive. Finally, to further validate the accuracy of the model, we conduct new simulations on polymers of various sizes, and investigate the role of jamming of the crowders.

## 1 Introduction

The mechanics underlying the conformation of bacterial DNA and its functions involve the physics of polymer collapse.^1–3^ In *Escherichia coli*, a more than a millimeter long genomic DNA occupies a region of less than a *μ*m^3^ called the nucleoid.^4^ Entropic crowding, caused by the depletion effect of small neutral molecules like polyethylene glycol on the DNA as a polymer, has been shown to be necessary in explaining the formation of a compact nucleoid *in vitro*.^5–8^ *In vivo*, osmotic shock and mechanical cell-size perturbations confirm this picture,^9^ and show that a 30% increase in the crowders (ribosomes and proteins) concentration can lead to a 3-fold decrease in nucleoid size, while association of DNA-binding proteins also clearly plays a role in this process^10^.

To explain these observations, multiple simulation studies have explored the impact of neutral crowding agents on a polymer, which is represented as a chain of beads.^1,9,11–13^ Since explicitly accounting for the crowder dynamics is computationally expensive, particularly when numerous small crowders are required, some studies have described the presence of crowders as an effective short-range attraction.^1,14^ These theories typically leverage the classical framework of the Asakura-Oosawa theory for depletion forces^15,16^ to describe the effective attraction between the beads, and to estimate its strength as a function of the size and density of the crowders.

Other numerical investigations explicitly accounted for the presence of crowders, represented by a fluid of hard spheres,^11–13^ to study their direct impact on polymer folding. In all of these studies, the authors simulated the collapse of a polymer comprising of *N* monomers, of fixed monomer size (which we call *D*), and with a mean distance between adjacent monomers *b* ≡ *D*, in the presence of crowders with a diameter *d* and a crowder volume fraction *ϕ*. All these studies concur that an increase in crowding results in a decrease in solvent quality. To demonstrate this, they examine quantities related to the swelling ratio of the chain, denoted as *α* = *R*(*ϕ*)*/R*(*θ*). Here, *α* represents the ratio between the end-to-end distance of the polymer chain under crowding conditions and its value in ideal conditions, where repulsive and attractive effects between monomers are balanced, and the polymer behaves as a ghost chain. The behavior of *α* as a function of *ϕ* reveals a sharp decline, indicating a polymer coil-to-globule transition, which occurs at different values of *ϕ* for different *d*.

To compare the varying conclusions, let us discuss these studies in further detail. Shendruck and coworkers^11,17^ concluded that the entropic crowding interaction is sufficient to trigger a second-order coil-to-globule transition, but that the Asakura-Oosawa theory was not sufficient to predict the form of the depletion pair potential. They proposed, as a solution, the use of a multi-parameter morphometric thermodynamic approach for the pair potential. Kang *et al*.,^12^ instead, observed in their simulations that the polymer collapse may or may not reach a globular phase. The behavior of the polymer is determined by a dimensionless control parameter defined as *x*_*Kang*_ = *R*(0)*ϕ* ^1/3^*/d*. We will refer to this as the “Kang parameter” in the following. Its value is derived from scaling argument and controls the statistical behavior of the polymer with respect to the crowing ratio. Finally, Jeon *et al*.,^13^ argued that their simulations supported the thesis that the swelling of the chain was a unique function of a different dimensionless control parameter defined as *x*_*Ha*_ = *ϕ D/d* (which we will call the “Ha parameter” in the following). This order parameter differs from the one proposed by Kang and coworkers. They argued, using approximations, that this parameter is compatible with the Asakura-Oosawa theory, in contrast to other studies that concluded the Asakura-Oosawa interactions alone cannot fully account for their results. Furthermore, they observed that the transition from coil to globule universally occurs for values of *x*_*Ha*_ that are close to 1. In brief, while these three different studies employed the same type of simulations, yielding consistent quantitative results, they accessed different regimes and analyzed the data using different paradigms, affecting their interpretations.

Here, we conduct an integrated analysis of the simulation results from these studies, aiming to capture the unifying traits. These traits lead us to propose a straightforward and effective mean-field theory capable of accurately describing all the simulation results and extend it to various scenarios. Moreover, to validate our predictions in novel regimes and examine the impact of jamming of the crowders, we conduct our own molecular simulations of polymer chains with explicit crowders.

## 2 Methods

### 2.1 The Flory mean-field theory for real polymers

We start by a short introduction of a Flory-like mean-field theory, which is the basic ingredient of our approach. In polymer physics, a chain of non-interacting monomers is described as a random walk, often called an “ideal chain”. In this simple model, the mean square end-to-end distance of the chain, denoted as ⟨*R*^2^⟩, can be expressed as

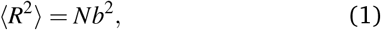

where *N* is the number of monomers in the chain and *b* represents the mean distance between adjacent monomers.

The classic Flory theory^18,19^ starts from this model and aims to predict the mean end-to-end distance of a “real chain” with interactions as the minimum of a free energy function. For the ideal chain, this free energy can be formulated as follows

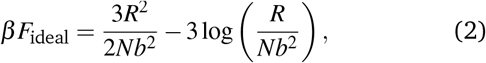

where *β* = (*k*_*B*_*T*)^−1^ represents the inverse temperature, with *k*_*B*_ the Boltzmann constant, and *T* the temperature.

When considering an interacting chain, we can introduce the swelling ratio *α* in the theory, which represents the ratio between the root mean square end-to-end distance *R* of the real chain and the one of the corresponding ideal chain, which is given by the expression *α* = *R/*(*bN*^1/2^). Consequently, the free energy of the ideal chain reads

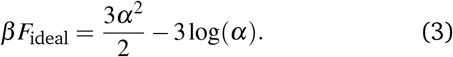

To obtain an expression for the free energy of the real chain, the interactions between the monomers and the solvent, as well as the interactions between the monomers themselves can be analyzed separately. The total free energy, *F*_tot_ = *F*_ideal_ + *F*_interactions_, is then the sum of the ideal chain free energy, *F*_ideal_, and the free energy contribution due to the interactions, *F*_interactions_, which can be described using a virial expansion, in a similar way as done in the theory of real gases,^20^

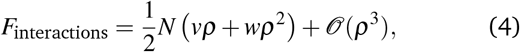

where *ρ* = *kN/R*^3^ represents the monomer number density with a geometrical prefactor *k* = 3/4π. The parameters *v* and *w* correspond to 2-body and 3-body monomer-interaction terms, respectively. When there is no attraction between monomers, these terms account for the steric repulsion between beads, and they reduce to *v*_0_ = 4*πD*^3^/3 (the monomer volume) and 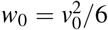. The 3-body interaction term *w* is taken as *w*_0_, and the total free energy can then be expressed as

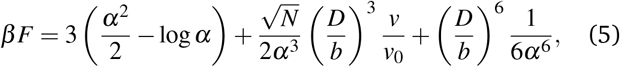

where the swelling ratio *α* can be obtained by minimizing the free energy, i.e. setting *dF/dα* = 0. This leads to the equation of state of the Flory mean-field theory for real polymers, which was derived by De Gennes in 1975^21,22^

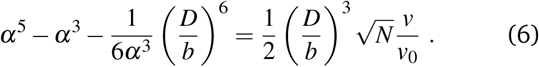

We use this equation to build our model. This equation is solved using the fsolve function from the python Scipy package. Further, in Sec. 3.4 the sets of analytical equation are numerically solved using the `minimize` function from pythin Scipy package, with Nelder-Mead algorithm (non-linear simplex).

### 2.2 Swelling ratio for the published simulation datasets

We perform a detailed analysis of the simulations carried out by Shendruck^11^ and Jeon *et al*.^13^ The details of the Shendruck *et al*. dataset^11^ are presented in Table 1. The observable used in their plots is the radius of gyration, denoted as *R*_*g*_. In order to obtain the swelling ratio, we first normalize the radius of gyration with its value in absence of crowders, *α*_0_ = *R*_*g*_(*ϕ*)*/R*_*g*_(*ϕ* = 0). The difference between *α*_0_ and the swelling ratio *α* lies in the denominator. Therefore, *α* is proportional to *α*_0_, with a coefficient of proportionality that is independent of *ϕ* and *d/D*, but dependent on the polymer length *N* and the ratio between the mean distance of adjacent monomers and the monomer size *b/D* ≡ 1. This relation will be used throughout the whole article. This coefficient can be determined as the value of *α* in the absence of crowders (where *ϕ* = 0, *α*_0_ = 1, and *v/v*_0_ = 1). We obtain its value by solving Eq. 6 with *v/v*_0_ = 1, which leads to the following relation

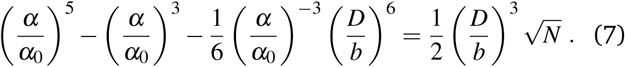

This equation allows us to express the data from Shendruck and coworkers in terms of the swelling ratio *α*.

**Table 1.** List of simulation data used in this work. Each row indicates the reference from which the simulation data was taken, the number of monomers *N*, the size ratio *D/d* between monomers and crowders and the range of crowder packing fraction values spanned in each work. In all cases, the chains and crowders are modeled as described in Sec. 2.3

| Ref. | $N$ | $\frac{D}{d}$ | $\phi$ |
| --- | --- | --- | --- |
| Shendruck et al.<sup>11</sup> | 15 | 3.00 | < 0.45 |
| Shendruck et al.<sup>11</sup> | 15 | 4.00 | < 0.45 |
| Shendruck et al.<sup>11</sup> | 15 | 5.00 | < 0.45 |
| Jeon et al.<sup>13</sup> | 50 | 2.00 | < 0.50 |
| Jeon et al.<sup>13</sup> | 50 | 2.50 | < 0.50 |
| Jeon et al.<sup>13</sup> | 50 | 3.33 | < 0.50 |
| This paper | 60 | 5.00 | < 0.60 |
| This paper | 60 | 2.50 | < 0.60 |
| This paper | 60 | 1.67 | < 0.60 |
| This paper | 60 | 1.25 | < 0.60 |
| This paper | 120 | 5.00 | < 0.60 |
| This paper | 120 | 2.50 | < 0.60 |
| This paper | 120 | 1.67 | < 0.60 |
| This paper | 120 | 1.25 | < 0.60 |

The details of the dataset by Jeon *et al*.^13^ are also summarized in Table 1. In their study, the measured observable is the normalized radius of gyration *α*_0_. To determine the swelling ratio *α*, we employ the same approach as used for the Shendruck dataset.

### 2.3 Molecular simulations of chains with explicit crowders

Here, we provide a brief overview of the model and molecular simulation method used to generate additional simulation data on the collapse of chains in the presence of explicit crowders. To ensure consistency, we adopted similar modeling choices as described in Refs. 11,13. Specifically, the DNA is represented as a linear chain comprising *N* spherical beads of diameter *D*. On the other hand, the crowders are modeled as spheres with a diameter of *d*. The interactions between all sites, including monomer-monomer, crowder-crowder, and monomer-crowder interactions, are described using a truncated and shifted Lennard-Jones potential, commonly known as the Weeks-Chandler-Andersen (WCA) potential.^23^ This purely repulsive pair potential is expressed as follows

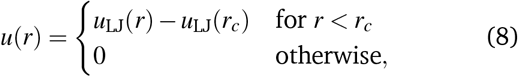

where *u*_LJ_(*r*) represents the Lennard-Jones potential given by

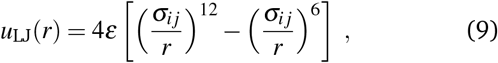

where *r* is the distance between the centers of the two beads, and *ε* and *σ*_*i j*_ are the energy and length scales of the interaction, respectively, with *i, j* either a crowder *c* or a monomer *m*. For our system, we define *σ*_mm_ ≡*D, σ*_cc_ ≡ *d*, and *σ*_mc_ = (*σ*_mm_ + *σ*_cc_)/2. The cut-off radius was set to *r*_*c*_ = 2^1/6^*σ*_*i j*_ to ensure a smooth and purely repulsive potential.

Adjacent monomers in a polymer chain are connected by the Finite Extensible Nonlinear Elastic (FENE) potential,^24^ which is given by

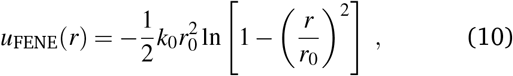

where *k*_0_ represents the spring constant, and *r*_0_ is the range of the potential. In our case, we employed the conventional values for polymers, specifically *k*_0_ = 30*ε/D*^2^ and *r*_0_ = 1.5*D*.^25,26^ This potential is purely attractive and competes with the purely repulsive WCA potential. The equilibrium point between these two potentials occurs at *r* ≈ 0.96*D*, which justifies our assumption that *b/D* = 1.

We perform Molecular Dynamics (MD) simulations in the canonical ensemble with a fixed number of beads (monomers and crowders) *N*_t_ = *N* + *N*_c_, volume *V* and temperature *T*. The mass of the monomers and crowders was considered as the unit of mass *m*, while the units of length, energy, and time are *D, ε* and 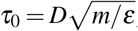 respectively. The equations of motion were integrated using the velocity Verlet method with a timestep *δt* = 0.005*τ*_0_. To maintain the desired temperature of *T* = 1.0*ε/k*_*B*_, we implement a Langevin thermostat with a damping constant of 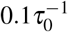. All simulations were performed in cubic boxes with a side length of *L* = 3*R*_*g*_(*ϕ* = 0), where *R*_*g*_(*ϕ* = 0) represents the radius of gyration of the chain in the absence of crowders. This choice of box size, which depends on the polymer length, was made to avoid artificial finite-size effects during the MD simulations. Additionally, we employ three-dimensional periodic boundary conditions. We perform simulations of single chains of length *N* = 60 and 120 immersed in solutions where the volume fraction of the crowders was set by the *x*_*Ha*_ = *ϕ D/d* parameter. In particular, we study systems where 0.15 ≤*x*_Ha_ ≤ 1.50, for size ratios between crowders and monomers of *d/D* = [0.2, 0.4, 0.6, 0.8]. Unless specified otherwise, the total number of MD steps of the simulations is 5 ×10^7^, and we collect statistics on the radius of gyration and end-to-end distance of the chains every 1000 steps. All the MD simulations are performed using the software package LAMMPS.^27^

## 3 Results

### 3.1 Previously proposed order parameters do not capture crowder-induced polymer collapse from different simulations

The size of a polymer under the effect of molecular crowding inherently depends on a wide range of variables that collectively determine its macroscopic state (Fig. 1A). If a unique dimensionless order parameter exists, derived from all these variables, this parameter unambiguously defines the size of the polymer, significantly simplifying the problem of predicting the compaction state of the polymer. In such a scenario, all the swelling ratios would fall on a single master curve when plotted as a function of this dimension-less order parameter.

**Figure 1.**
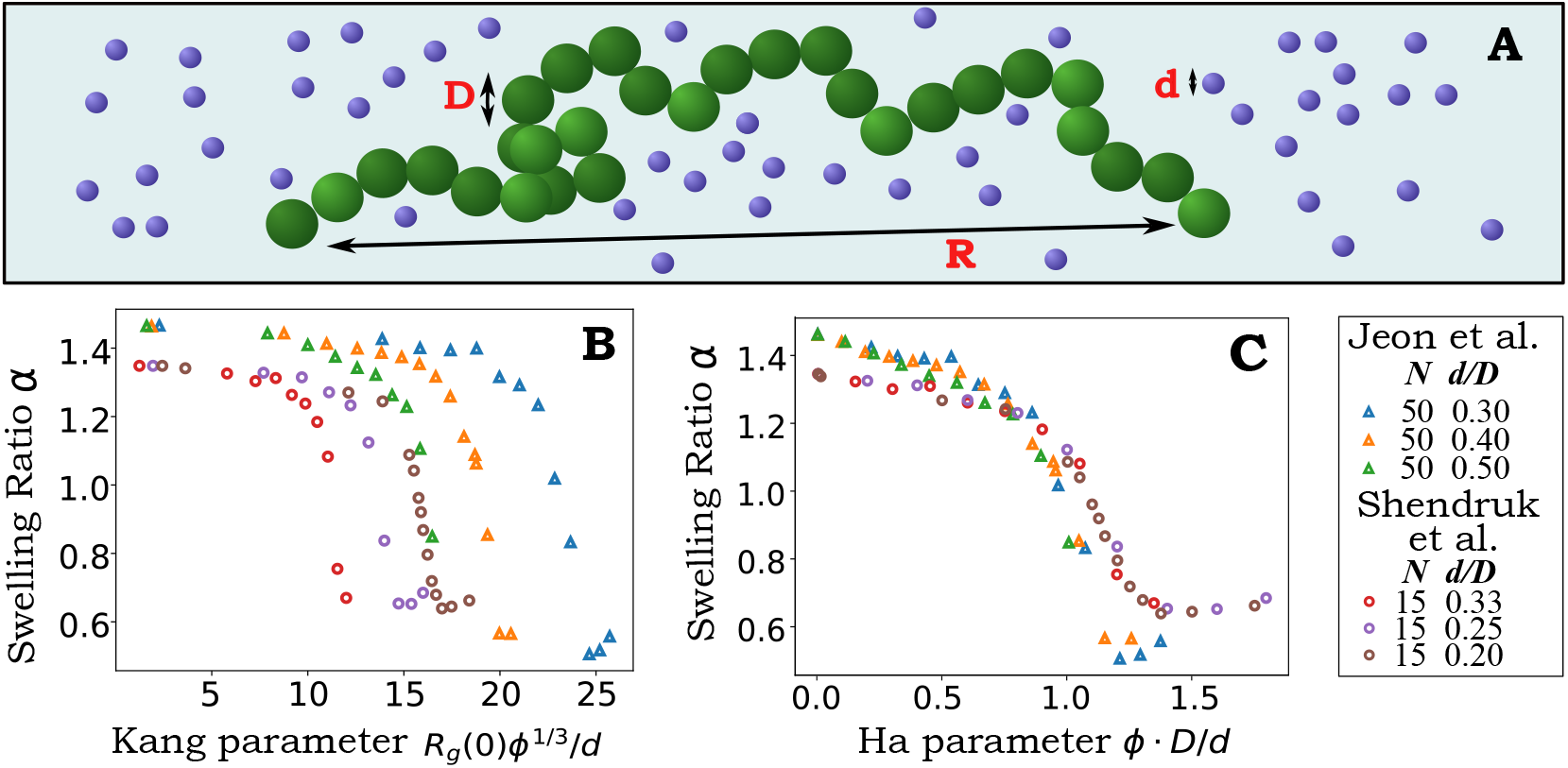
Joint analysis of the swelling ratio *α* in crowder-induced single polymer collapse from different simulation studies. **(A)** Sketch of a bead-spring polymer in a crowded medium of hard spheres, illustrating the main simulation parameters common to all the studies considered here. Monomers have size *D* and crowders have size *d* and volume fraction *ϕ*. **(B)** Swelling ratio *α* (simulations from Refs. 11,13), plotted as a function of the parameter *x*_*Kang*_ proposed by Kang and coworkers.^12^ **(C)** Same data, plotted as a function of the control parameter *x*_*Ha*_ proposed in Refs. 13,28.

Following this line of reasoning, we collected all the swelling ratios from the simulation data of Shendruck *et al*.^11^ and Jeon *et al*.^13^ (Table 1) and plotted them as a function of the two order parameters *x*_*Kang*_ = *ϕ* ^1/3^*R*(0)*/d* or *x*_*Ha*_ = *ϕ D/d* as identified in these studies (see Introduction). Fig. 1 demonstrates that both the Kang parameter *x*_*Kang*_ (Fig.1B) and the Ha parameter *x*_*Ha*_ (Fig. 1C) are insufficient to fully describe the swelling ratio of both datasets. In each plot, the swelling ratios from different simulations appear on distinct curves when plotted against the corresponding order parameter. This observation indicates that neither of these order parameters alone can capture the complexity of the polymer collapse under molecular crowding conditions.

On the other hand, we observe that the swelling ratios of polymers with the same chain length *N* collapse onto a single curve when plotted as a function of *x*_*Ha*_. However, when comparing polymers of different lengths, each of them falls on a different curve. This indicates that the Ha parameter alone is not sufficient to account for the variation in swelling behavior among polymers with different chain lengths. Thus, based on these observations, we conclude that the simulations conducted by Jeon and coworkers, which solely focused on the *N* = 50 polymer length, did not fully account for the role of chain length *N* on the swelling ratio of crowder-induced polymer collapse.

Moreover, both at low concentrations of crowders and under conditions where crowding induces polymer collapse, the radius of gyration exhibits a clear dependence on the chain length *N*. Therefore, we conclude that the chain length appears to be an essential factor, which needs to be considered to accurately predict the behavior of the polymer collapse in the presence of crowders.

This *N* dependency is well described by standard polymer theory, which predicts that the radius of gyration scales as *R*_*g*_ ∝ *N*^3/5^ for a polymer in a good solvent, *R*_*g*_ ∝ *N*^1/2^ at the transition point, and *R*_*g*_ ∝ *N*^1/3^ for the globular state^18,29,30^ at a mean-field level. These theoretical predictions provide a solid basis for understanding the observed variation in the radius of gyration with respect to the chain length *N* in different crowding conditions. In terms of the swelling ratio, this implies that *α*(*ϕ* = 0) ∝ *N*^1/10^ and *α*(*ϕ* → 1) ∝ *N*^−1/6^.

To sum up, we can confidently rule out the existence of a single parameter such as *x*_*Ha*_ or *x*_*Kang*_. The reason is that no single rescaling of the crowders’ volume fraction can account for the different scaling in polymer length *N* observed in the swollen phase compared to the globular phase.

### 3.2 The second-order virial coefficient is approximately set by *x*_*Ha*_ = *ϕ D/d* as the relevant dimensionless parameter

In light of the results from the previous section, we explore the possibility of finding an intensive quantity (independent of the polymer size *N*) that could be, at least approximately, described by a single dimensionless parameter. To achieve this, we turn to the Flory mean-field theory,^21,22^ which enables us to extract the quality of the solvent, represented by the second virial coefficient *v*, from the swelling ratios (see Methods).

More specifically, the Flory mean-field theory establishes a relationship between the polymer size *N*, the swelling ratio *α*, and the second virial coefficient *v* through the following equation of state

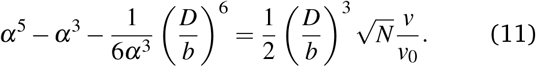

For a detailed derivation of this equation, see the Methods section 2.1. This equation provides a means for quantifying the quality of the solvent based on the swelling ratios of the polymer and the second-order virial coefficient. The first two terms in the equation arise from the free energy of the ideal chain, while the third term and the right-hand side account for the interactions between monomers in the presence of a solvent composed of neutral crowders. This formula takes into account solely second- and third-order terms in the virial expansion. Both of these terms are independent of *N* and can be treated as unspecified functions of *D, d*, and *ϕ* when the solvent solely consists of neutral crowders.

It is important to note that three-body interactions can be neglected for a polymer in a good solvent. Therefore, this term becomes only relevant when the polymer undergoes a transition to a globular form in a very bad solvent. Furthermore, higher-order interactions are also expected to be negligible. While the derivation of Eq.11 is not strictly rigorous (as long-range correlations were shown to be important for a polymer chain^29,31^), it is often considered to be a good approximation for reasons that are more generic than its derivation.^32^ Specifically, Eq. 11 conveniently combines three asymptotic behaviors, corresponding to different universality classes, into one compact phenomenological formulation: *R*_*g*_ ∝ *N*^1/2^, representing the ideal chain behavior observed near the collapse transition (*v* ≈ 0), *R*_*g*_ ∝ *N*^3/5^, describing the mean end-to-end distance *R* behavior of a self-avoiding chain in a good solvent (*v* ≫ 0), and *R*_*g*_ ∝ *N*^1/3^, characterizing the globule state in a bad solvent condition (*v* ≪ 0). Hence, Eq. 11 serves as an interpolation formula able to capture these three distinct scaling behaviors.

Figure 2 shows that all the *v/v*_0_ values extracted from the swelling ratios in simulations using Eq. 11 are well-defined by the Ha parameter *x*_*Ha*_ = *ϕ D/d*, nearly collapsing onto a single curve. Remarkably, the discrepancies observed in the asymptotic scaling for different values of *N*, as seen in the plot of *α* versus *x*_*Ha*_ (Figure1C), are successfully resolved when plotting *v/v*_0_ versus *x*_*Ha*_.

**Figure 2.**
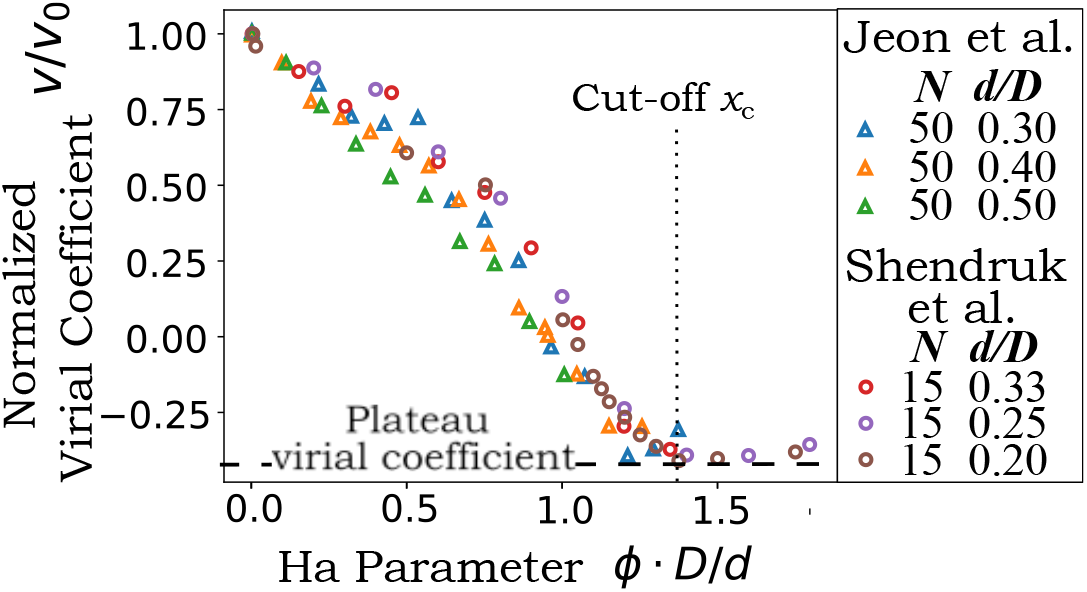
Near collapse of the second-order virial coefficients extracted from published simulation data (Refs. 11,13) plotted as a function of *x*_*Ha*_ = *ϕ D/d*, the parameter proposed by Ha and coworkers.^13,28^ The plot supports a good collapse across the coilglobule transition, with some discrepancy for intermediate values of *x*_*Ha*_.

The value of *v/v*_0_, by definition, was set to unity when *x*_*Ha*_ → 0 (see Methods), and we find that it levels off to a constant value, as identified from the simulated data as (*v/v*_0_)_*c*_ ≃−0.35 for *x*_*Ha*_ greater than *x*_*c*_ ≃ 1.25. This value is independent of the simulation parameters considered, suggesting a robust and universal behavior for (*v/v*_0_)_*c*_ and *x*_*c*_.

In conclusion, the collapse of all data points onto a single curve is highly satisfactory, although not entirely perfect. It is also essential to acknowledge that these numerical values for (*v/v*_0_)_*c*_ and *x*_*c*_ may not be entirely universal and could depend on specific details in the potentials defining the simulated polymers and crowders. However, no discrepancies were observed in any of the available simulations.

### 3.3 Derivation of an effective potential for the implicit solvent

Notably, the “inverse” analysis presented in Fig. 2 enables the development of an effective “direct” mean-field theory.

As previously mentioned in the Introduction, multiple studies concluded that the Asakura-Oosawa theory for colloidal spheres in the presence of depletants is not directly applicable to the collapse of a polymer in the presence of crowders. In the Asakura-Oosawa theory,^15,16^ entropic crowding is effectively incorporated through an attractive pairwise short-range potential. Assuming that a different effective short-range potential can describe our system, we have derived a simple formula for this effective potential energy between two monomers. Specifically, we considered a square-well potential *U* (*r*) with a range of *D* + *d* and a hard-core repulsion of size *D*, as illustrated in Fig. 3A. The advantage of this simple discontinuous potential lies in its dependence on just one parameter, *β E*_0_, representing the depth of the potential.

**Figure 3.**
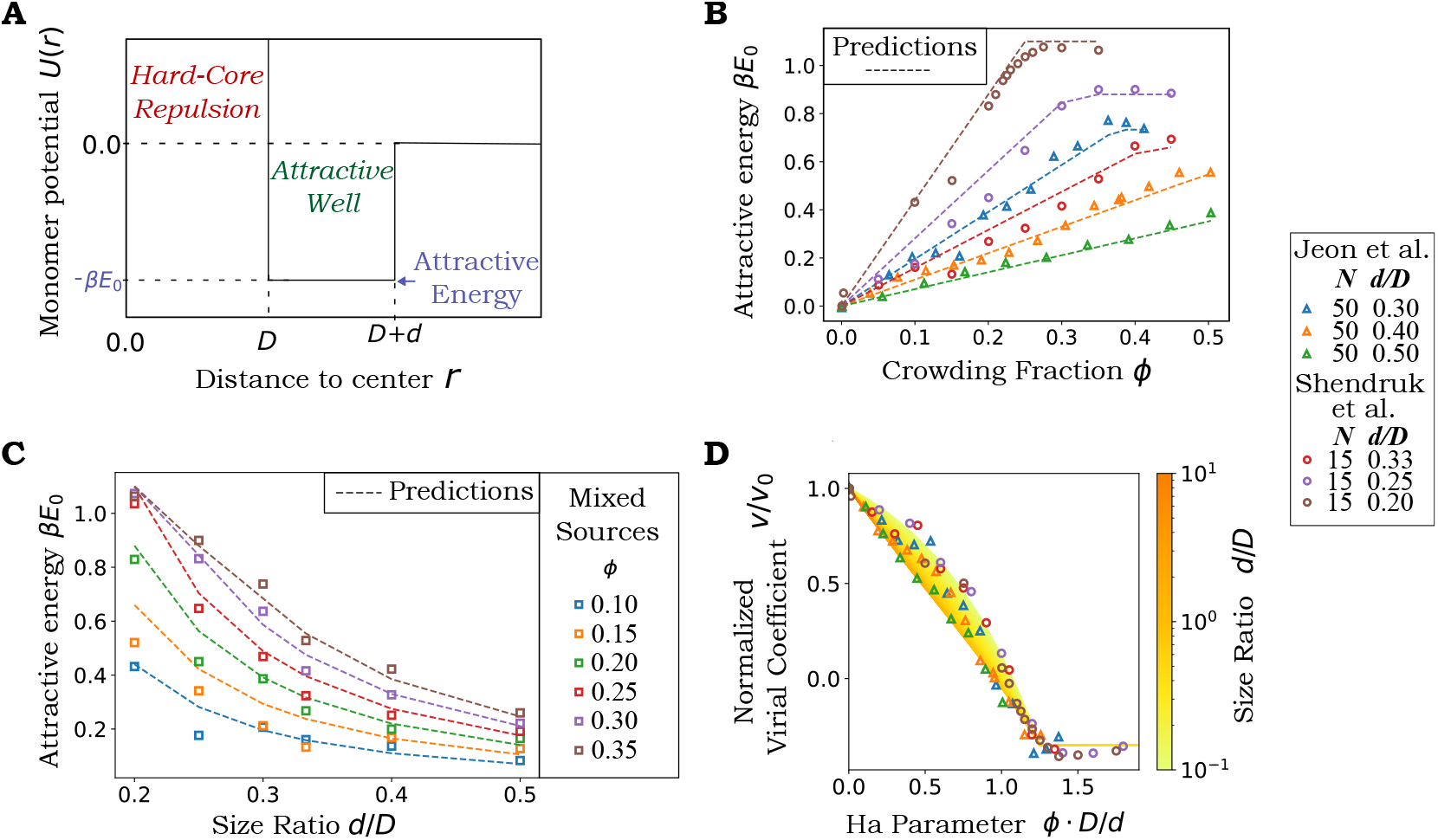
A Flory-like mean-field model captures the crowder-induced coil-globule transition. **(A)** The model approximates the interaction between monomers as the sum of a hard-core repulsion and a monomer-monomer square-well attraction with a range equal to the crowder size *d*. **(B)** Depth of the effective energy well *β E*_0_, plotted as a function of crowder packing fraction *ϕ*. Triangles represent simulation data for different values of relative crowder size *d/D*, and the dashed lines are fits of the empirical potential derived in the main text (Eq. 15). **(C)** Depth of the effective energy well *β E*_0_, plotted as a function of relative crowder size *d/D*, simulation data (squares) are interpolated from the curves on panel B. **(D)** Normalized second virial coefficient *v/v*_0_ as a function of the Ha parameter. The data are the same as in Fig. 2, theory (color-coded solid lines) highlight the increasing non-linear effects as a function of *d/D* (obtained by combining Eqs. 12 and 15), which predict the imperfect collapse observed in the simulation data.

From a fundamental perspective, the specific shape of the potential is expected to be of minor importance as long as it remains short-ranged. Therefore, we assume that the range of interactions corresponds to the size of the crowding agent *d*. This choice allows us to capture the essential features of the interactions between the monomers and crowders in a simple yet effective manner. The functional form for *E*_0_ = *E*_0_(*ϕ, d/D*) is derived from the second virial coefficient:^33^

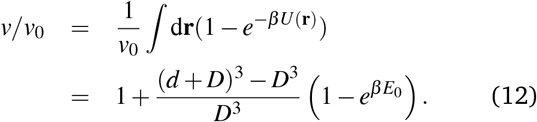

We note that if we assume that the virial coefficient collapses with *x*_*Ha*_, the interaction energy *E*_0_ cannot be expressed solely as a function of a single parameter *x*_*Ha*_. Instead, it needs to be represented as a function of both *d/D* and *ϕ*. From the plot of *E*_0_, calculated from *v/v*_0_ through Eq.12, as a function of *ϕ* (Fig.3B), we observe that *E*_0_ can be approximated as a linear function of *ϕ* with a plateau at the coil-globule transition, occurring when *x*_*Ha*_ = *x*_*c*_. Within the range of parameters considered in the simulations, this simple observation holds true. By definition, at the transition to the plateau, the value of the second-order coefficient becomes *v/v*_0_ = (*v/v*_0_)_*c*_.

Based on this simple observation, we can calculate the first-order approximation in *d/D* for the value of this plateau, denoted as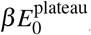, which can be expressed as

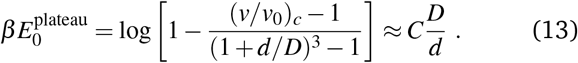

By fitting Eq. 13 to the simulated data, we determine the constant *C* to be 0.223 ± 0.003. The numerical value of the constant *C* measured from the available simulations, combined with the fact that in the absence of crowders there is no attractive interaction (*E*_0_ = 0), suggests the following simple potential as a model,

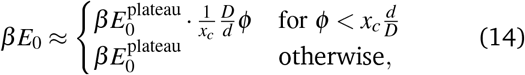

and its first-order approximation on the right hand side of Eq. 13 is,

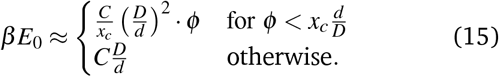

Comparing this approximation to available simulations, we see that it works very well in the range of parameters observed (0.20 < *d/D* < 0.50), since it agrees with the data for both *E*_0_ as a function of *ϕ* for different *d/D* (Fig. 3B, dashed lines) and *E*_0_ as a function of *d/D* for different *ϕ* (Fig. 3C, dashed lines). Interestingly, this result corresponds to a scaling of the interaction energy 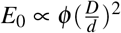that does not correspond to the classic Asakura-Oosawa energy scaling (see Appendix B for a rationalization of why this is the case), which confirms previous observations.

Finally, considering that *E*_0_ and *v/v*_0_ are linked by Eq. 12 and that the interaction energy *E*_0_ depends on *d/D* and *ϕ* separately, the model predicts that the empirical near-collapse of *v/v*_0_ by a single dimensionless parameter *x*_*Ha*_ reproduced in the previous Section is, in fact, not universal. We should instead consider this collapse as the simplest empirical approximation that is valid for the range of parameters considered in the available simulations. In fact, by numerically combining Eq. 12 with Eq. 14, the model gives a different approximation, which gives a richer and more precise description of the observed data for *v/v*_0_ as a function of *x*_*Ha*_ (Fig. 3D). Note that our prediction is not a single universal function of *x*_*Ha*_, but the mean-field model predicts a gradient of shapes for different crowder sizes *d/D*, as, shown in Fig. 3D. The same figure shows that the predicted gradient reproduces a trend that is visible in the simulated data sets.

### 3.4 The identified effective potential generates correct predictions for simulations of confined polymers and ring polymers

So far, we have focused on the case of a linear unconfined polymer. However, as mentioned in the Introduction, motivated by the organization of bacterial genomes, simulations in the literature also explored scenarios involving ring polymers and/or polymers in confinement. In light of this, we pose the question of whether our effective model could be generalized straightforwardly to encompass these situations. These modifications influence the overall free energy landscape of the polymer, but should only marginally change the monomer-monomer interactions (which are represented in our model as local interactions). The main correction to the current model would arise from surface adsorption on the confining edges, which we neglect here for simplicity. Under this assumption, the effective pairwise interaction energy that characterizes our model should also be valid for ring-shaped and/or confined polymers, and only the Flory formula from De Gennes (Eq. 11) should be modified. Since ring polymers tend to adopt a linear two-strand configuration,^35^, and the diameter occupied by a ring polymer corresponds approximately to the one occupied by a linear polymer of half its length^36^, the ring polymer free energy can be approximated as a chain polymer with half the number of monomers. Hence 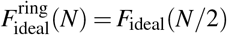; in other words, the volume of a ring polymer is approximately equal to the one of a linear chain with half the number of beads.

We can write:

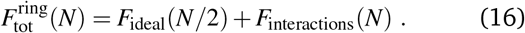

We adopted this simple approximation and compared our prediction with numerical simulations. Chauhan *et al*.^34^ simulated a ring polymer in crowded environment. Fig. 4A compares their simulation data (triangles) to the predictions of our modified model. The comparison (which involves no free parameters) shows an excellent qualitative agreement and a satisfactory quantitative agreement.

**Figure 4.**
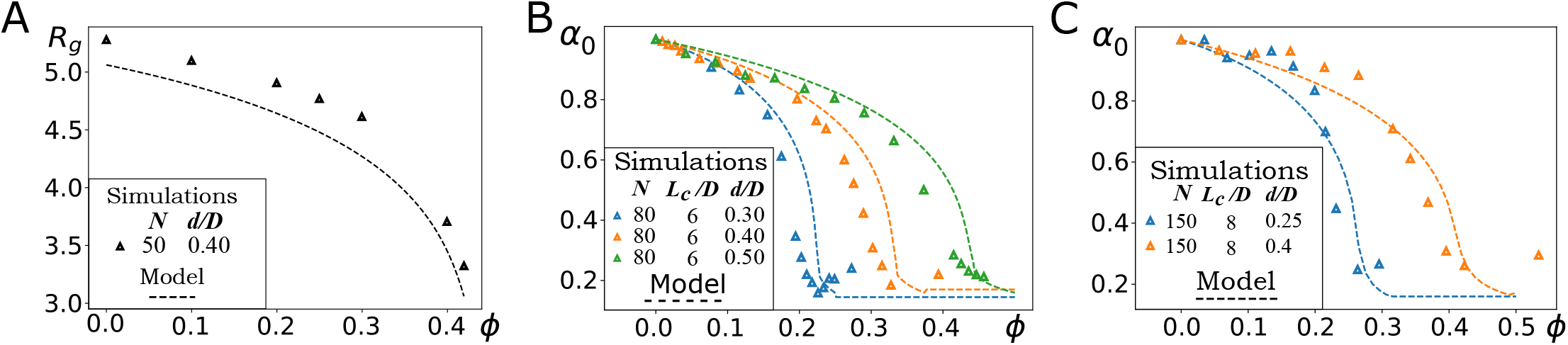
Simple modifications of our model make effective predictions for a confined linear polymer, a free ring polymer and a confined ring polymer. **(A)** Prediction of the collapse curve, *R*_*g*_ (in units of *D*) versus *ϕ*, for a confined linear polymer compared with simulation data from Ref. 28. **(B)** Prediction of the collapse curve, *α*_0_ versus *ϕ*, for a free ring polymer compared with simulation data from Ref.34. **(C)** Prediction of the collapse curve, *α*_0_ versus *ϕ*, for a confined ring polymer, compared with simulation data from Ref. 9. *α*_0_ is defined as *R*(*ϕ*)*/R*(*ϕ* = 0). The above comparisons involve no free or adjusted parameters, as all parameters in the model are fixed from simulation input parameters.

To describe a confined polymer, we extend our model with the classic blob model from Pincus,^37^ for a cylindrical confinement of diameter *L*. Specifically, we used the non-confined free energy to compute the number of monomers *N*_*L*_ that a free chain would need to have a size *R*(*N*_*L*_) = *L*. Subsequently, when *N* < *N*_*L*_, we employ the free energy of a free chain; conversely, for *N* ≥ *N*_*L*_, we describe the polymer as a chain of *N*_*c*_ = *N/N*_*L*_ confinement blobs. In this case, the overall end-to-end distance becomes the product of the number of confinement blobs and the diameter of each blob. Hence, the confinement diameter reads *R*(*N*) = *N*_*c*_*L*. This straightforward modification of our model allows us to formulate another prediction, which we can compare with simulations. Kim *et al*.^28^ simulated polymers in cylindrical confinement with explicit crowding. Fig. 4B compares the simulation data (triangles) to the predictions of our model, once again finding good agreement. Finally, Yang *et al*.^9^ simulated ring polymers in cylindrical confinement subjected to crowding. In order to describe this situation, we combined the two above modifications, finding again a good match between the predictions of our model and the simulation data, with no adjustable parameters (Fig. 4C). These results confirm the validity of the phenomenological form of the effective interaction energy as a function of the volume fraction of the crowders. Moreover, they underscore its adaptability for different applications.

### 3.5 Polymer collapse does not take place for crowders above a critical size, as predicted by the theory

As described in Section 3.3, in all the simulations considered in this study the polymer reaches the collapsed state when the densities of crowders are above the critical value defined by 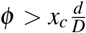. It is evident that due to its definition (*ϕ* ≤ 1), there exists a physical limit on the size *d* of crowders that is capable of inducing the collapse of a polymer chain. However, simulations from Chanil et al. and Shendruck et al. where performed at values of *d/D* such that this limit wouldn’t be relevant and collapse would always be reached (Fig. 5A). To explore this limit, we conducted a fresh set of simulations (see Table 1) that enabled the examination of larger set of *d/D* values. By plotting the swelling parameter *α* against increasing *d/D* values, we can see how the polymer collapse transition shifts towards higher *ϕ* values. Notably, for crowder-to-monomer size ratios *d/D* ≥ 0.6, no collapse was observed within the range of *ϕ* values investigated in our simulations (see Fig. 5B). Particularly at *ϕ* values near the point where hard spheres cease to exhibit fluid-like behavior, the collapse was hindered by a slowing down in the dynamics of the crowders. From these observations, we conclude that the collapse transition can be achieved in simulations when *ϕ* ≤ *ϕ*_*c*_, where *ϕ*_*c*_ ≈ 0.6. This value is lower than the theoretical limit of 1 and closer to the random close-packing density.^38^ Consequently, this establishes a threshold for the size of crowders capable of inducing polymer collapse (*d/D* < *ϕ*_*c*_*/x*_*c*_), while for larger *d/D* ratios, this would result in a polymer immersed in a jammed solvent.

**Figure 5.**
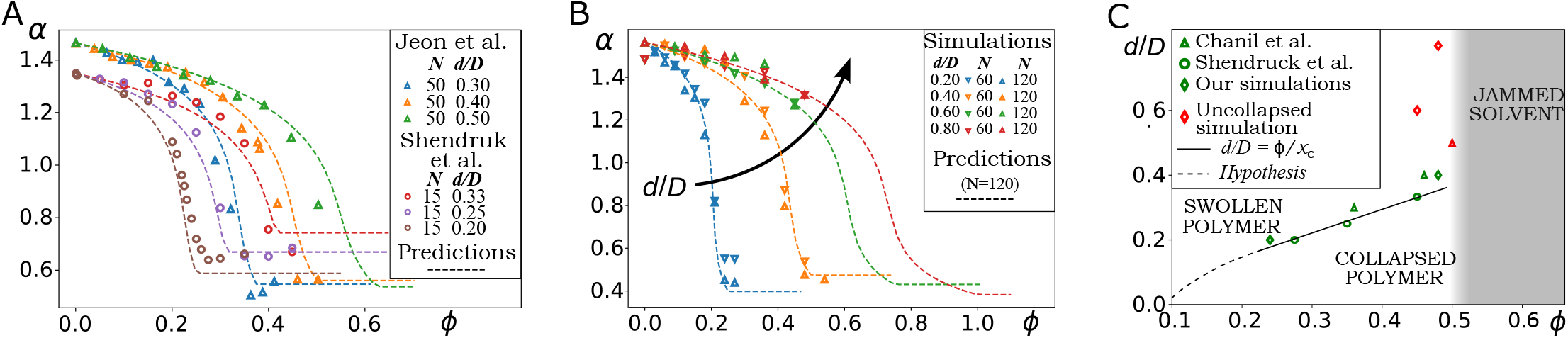
Polymer collapse can be prevented by jamming of the crowders for large volume fractions *ϕ* and size ratios *d/D*. **(A)** Simulations from Chanil et al.^13^ (triangles) and Shendruck et al.^11^ (circles). **(B)** Simulations for *N* = 60, 120 at different *d/D*. The arrow indicates that swelling ratios marginally increase with increasing *d/D*. For both panels the predictions of Eqs. 11, 12 and 15 are plotted in dashed lines (only for *N* = 120 in **(B)**). For high *d/D* and high *ϕ*, the transition point for the collapsed state is not reached and the prediction from our model no longer matches the simulations. **(C)** Phase diagram showing the value collapse values of *ϕ* (green markers). Collapse values are hand-picked at the value of *ϕ* where the swelling parameter curve flattens. For a selected subset of simulation parameters equilibration is not reached, we plot in red markers the maximum simulated *ϕ* for these cases. We gathered have data from Chanil et al.^13^ (triangles), Shendruck et al.^11^ (circles) and our simulations (diamonds). The solid line reports the solution for the transition between the collapsed and swollen phase according to Eq. 15, that is a prediction for the location of the green markers. The shaded area indicates the regime where the fluid phase of hard spheres becomes metastable with respect to the solid phase, this regime was not investigated with current simulations. The dashed line represents the hypothesis that the phase line should go to *d/D* = 0 as *ϕ* tends to 0.

The system under investigation has some analogies with the phase transition observed in 3D glass-forming supercooled^†^ binary mixtures,^39–43^ where jamming transitions occur at values of *ϕ*_*c*_ above 0.49. In binary mixtures, the larger spheres become glassy at lower densities compared to the smaller spheres. Thus, provided that the *d/D* ratio remains adequately small, the polymer made by these larger spheres undergoes a collapse before the crowders become jammed. In light of this existing body of literature, and drawing parallels with the proposal by Kang and colleagues^12^, we present a phase diagram for the polymer-crowders mixture as a function of *d/D* and *ϕ* (Fig. 5C). This phase diagram shows a division into three distinct regions with a triple point at (*ϕ, d/D*) = (*ϕ*_*c*_, *ϕ*_*c*_*/x*_*c*_). In one region, the polymer swells; in another, it collapses (potentially leading to jamming); and in the third, it remains unable to collapse due to solvent jamming or a phase transition to a solid state.

## 4 Discussion

In this work, we introduced a novel mean-field approach that employs a two-body effective interaction between monomers within a polymer immersed in a bath of crowders. Our approach assumes a square-well potential with a range that equals the size of the crowders. The Mayer function establishes a link between the depth of this potential and the second-order virial coefficient, which we extracted from simulations with explicit crowders. The resulting picture (see Fig. 3B-C) offers a direct model for the effective interaction, which smoothly interpolates between two distinct regimes: an “inelastic state” (the globule) where the short-range attractive energy remains constant with crowder density, and an “elastic state” (the coil) where the energy linearly increases with crowder density *ϕ*, starting at *E* = 0 for *ϕ* = 0. The density at which the polymer collapses is defined by the intersection of these two curves. Notably, as the second curve can be approximated linearly as shown by the data, our mean-field model naturally emerges, becoming entirely defined by these two regimes.

Depletion emerges as the underlying physical mechanism driving polymer collapse. However, unlike a first principle approach approach, our theory focuses solely on emergent properties. It achieves this by interpolating two distinct polymer phases and their specific mechanical characteristics, aligning with the spirit of the original Flory mean-field theory.

Our approach has the advantage of being both generic and simple, succeeding in areas where the classic Asakura-Oosawa theory does not fit quantitatively the phenomenology. As previously discussed,^11,12^ the Asakura-Oosawa potentials are formulated (and valid) solely in the regime of very low solvent concentrations *ϕ* < 0.2. However, none of the simulations available in the existing literature meet this criterion, as it significantly deviates from the “interesting” regime where polymer collapse can occur. Instead, many simulations are often executed in the proximity of the jamming transition for both the solvent and the polymer globular phase (see Fig. 5C). For this reason, we conjecture that a theory valid for jammed fluids might be appropiate for extending our mean-field observations.

Other approaches found in the existing literature for the considered regime are generally more complex and involve a higher number of parameters as compared to our model. These approaches include morphometric thermodynamics,^44,45^ 3rd-order virial expansions,^20^ and the scaled particle theory^46^. All of these approaches result in theories with a higher number of parameters than the Asakura-Oosawa theory, making interpretation more challenging and imposing more difficulty in estimating their limits of applicability.

Another interesting feature of our model is that it allows us to qualitatively reinterpret the phase diagram proposed by Kang and coworkers^12^ for the crowders-induced collapse of polymers, as we explained in Section 3.5. Kang and colleagues addressed the question of why molecular crowders do not significantly affect the conformation of disordered proteins^47,48^ but they affect the collapse of DNA molecules. Specifically, by testing the effect of neutral crowders on four intrinsically disordered polymer domains by single molecule fluorescence resonance energy transfer, Soranno et al. demonstrate that these domains can be affected either by a reduction of the end to end distance or by folding^47^. For the population of molecules that show a reduction in end-to-end distance, they tested the effects of crowder sizes and concentrations, and demonstrated that the effect is larger for larger crowder concentrations as well as for larger crowder sizes at fixed density, opposite to the scaling proposed by Ha and collaborators. The reduction in end-to-end distance is overall modest in all conditions, and its trend appears to change approximately linearly with crowder concentration. The authors conclude that in order to account for these effects one has to consider the interpenetration of the disordered domain and the polyethylene glycol constituting the crowder agent, which in our opinion applies only when crowder polymers are approximatively of the same size or larger than the diluted polymer on which this effect is measured. As such, we do not find it surprising that the Ha scaling and the simulations with hard spheres do not present the same trends. The conclusion drawn by Kang and coworkers is that proteins collapse only modestly since proteins are generally smaller than DNA molecules, and consequently they are less affected by crowding. Instead, our model (confirmed by simulations with hard spheres) suggests that it is not the size of the polymer, but the size of its monomers compared to that of the crowders that matters the most. Consequently, since DNA is made of large units (in the 10nm-100nm size range), it can be collapsed by protein-sized crowders due to depletion interactions. Since proteins are composed by amino acids having a contour length in the sub-nanometer range,^49^ this results in high *d/D*, and our phase diagram predicts that in these conditions jamming of the solvent prevents collapse. Hence, our model is not necessarily in contradiction with the conclusions of Kang and coworkers, since the Kang theory might become predictive when the finite-size effects and interpenetration of chains are the relevant phenomenon driving the end-to-end distance reduction, and it would be very interesting to test this hypothesis through numerical simulations. Alternatively, we propose here that an experiment could be designed where a large protein made of alternated disordered domains and weakly interacting folded domains would be complemented by crowders, since the large folded domains could effectively behave as large monomers linked by the disordered domains, changing the effective value of *d/D*. In this idealized case, our theory would predict that folded domains would assemble into a larger “globule”, depending on the levels and sizes of (neutral) crowding. In the theory by Kang and coworkers, instead, the presence of multiple folded domains would not change the behavior at fixed protein length. Hence, it might be possible to use this scenario as an experimental testable discriminant between the two theories, or, assuming both theories have a range of validity, to identify the crossover between the two theories by varying the size of the folded domains and the size of crowders. We speculate that the effects described by our theory will be predominant when crowders are slightly smaller then folded domains, and that jamming would be a predominant effect for larger crowders or smaller disordered domains, while the Kang theory may become predominant for crowders larger then the size of the protein in good solvent.

In general, our theory is currently limited by the fact that the monomer aspect ratio should be relevant, particularly for slender polymers like DNA, and we expect this to add an extra dimensionless variable to the problem. This will alter the numerical values of all the observed dimensionless constants extracted from the simulations and requires an extension to the theory proposed here. Similar ideas have been developed in Ref. 50. To fully elucidate this point, simulations of polymers with rod-shaped monomers or worm-like chains in explicit crowders will be required. While our theory is a step forward towards a quantitative descriptions of crowder-induced bacterial chromosome compaction,^2,6,9,13^ there are still many other aspects to address. First, it is based on simulations of polymers and embedding media that lack any biological activity. Chromosomes are found in a constant state of crowding where out-of-equilibrium effects that are relevant for polymers larger then the ones simulated so far could slow down the polymer collapse and lock it in a glassy state. Such states can result from topological barriers (*i. e*. knots)^51^ and pearling.^52,53^ Glassy states could theoretically survive for time scales comparable to the cell cycle rendering the theory of globular polymers inadequate to describe bacterial chromosomes. On the other hand, these out-of-equilibrium effects are mitigated by opposed active processes fueled by ATP, which can fluidize the dynamics of the cytoplasm^54,55^ and will be predominant over such timescales^56^; to a greater extent, active forces might play a fundamental role in nucleoid positioning^57^. Further, our model focuses only on the effect of monodisperse neutral crowders, while *in vivo* chromosomes are surrounded by both neutral and charged crowders, with a broad distribution of sizes, as well as relying on further interactions, e.g. from bridging proteins and active loop extruders.^2,58,59^. The size distribution of crowders turns the jamming effects fundamental to our theory into a scale-dependent phenomena: molecules with sizes equivalent to or larger than the dominant crowding agent will be more affected than small particles^60–62^. Finally, we have shown the validity of our theory for linear, confined and ring polymers simulations, while DNA in bacterial nucleoids is supercoiled and organised in topological domains such as plectonemes and toroids^63–66^. Even without considering crowders, simulations of supercoiled polymers show that this affects polymer structure and dynamics in many aspects (as reviewed in Junier et al.^67^). Further, it has been proposed that supercoiling can interact with crowding and affect the way this collapses the nucleoid^68,69^. A full theory or simulation encompassing the effects of crowding as well as a quantitative characterization of the effects of each of these elements on the compaction of bacterial nucleoids is still missing.

## Author Contributions

Conceptualization: MCL, VFS, QC. Data curation and analysis - model development: QC, VFS and MCL. Software-simulations: GG, GCV and MD. Supervision: MCL, VFS and MD. Writing - original draft: QC, VFS and MCL. Writing - review and editing: VFS, MCL and MD.

## Acknowledgements

We are grateful to Ignacio Pagonabarraga, Jean-François Joanny, and Daan Frenkel and Jaan Mannik for useful discussions. MCL was supported by AIRC - IG grant no. 23258.

## Appendix

### A Data analysis

Data were analyzed by custom-made Python jupyter notebooks with numpy, scipy, available at the following link: https://github.com/QuentinChab/PolymerCollapseWithCrowding

### B The Asakura Oosawa theory doesn’t fit the simulations

Asakura and Oosawa^15,16^ predicted the existence of an attractive short-range depletion force between two plates submerged in a bath of colloids. This force can be generalized for other geometries. The Asakura-Oosawa (AO) potential is in all cases equal to the osmotic pressure of the crowder’s gas multiplied by the gain in volume *δV* (*r*) accessible to this gas due to the proximity of the immersed bodies,

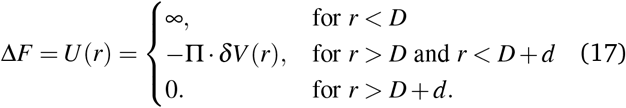

For spherical bodies, this volume consists of the overlap between two spherical shells,^70,71^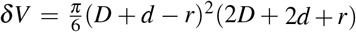.Eq.17 includes the infinite hard-core potential that models the hard spheres of diameter *D*, and is zero for distances greater than *r > D* + *d*. Hence, the AO theory predicts short ranged effective interactions. Because of this, Eq. 17 can be approximated by a square well potential^20^ of depth *E*_0_ delimited by the hard-core potential and the interaction range *D* + *d*, the same shape introduced by us in Section 3.3 and Fig. 3A. The value of *E*_0_ is commonly taken equal to *δV* (*r*) Π at *r* = *D*.^20,71^ As a first approximation, the osmotic pressure is taken linear with the crowders volume fraction 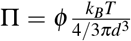, since in diluted conditions (*ϕ* < 0.3) the interactions between crowder particles can be neglected.^70,71^ This results in the following effective interaction energy,

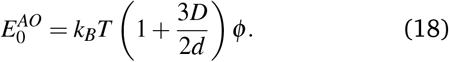

In the limit of small crowders (when *D/d* ≪ 1), this formula depends on *x*_*Ha*_ only, and this fact has been used in support of *x*_*Ha*_ being the key dimensionless parameter.^13^ However, Section 3.3 shows that in order to reproduce the simulation data, and thus the approximate scaling with *x*_*Ha*_, *E*_0_ should scale as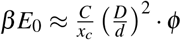. Hence, we conclude that the scaling relation predicted by this approximation of the Asakura-Oosawa theory (Eq. 18) is fundamentally not compatible with the relation between *D/d* and *E*_0_ obtained in simulations.

## Footnotes

† we mean here that in a simulation the system is prepared as a fluid and then undergoes structural relaxation and phase separation

